# Aspartate-phobia of thermophiles as a reaction to deleterious chemical transformations

**DOI:** 10.1101/2020.08.16.252783

**Authors:** Etienne Villain, Philippe Fort, Andrey V Kajava

## Abstract

Prokaryotes growing at high temperatures have a high proportion of charged residues in their proteins to stabilize their 3D structure. By mining 175 disparate bacterial and archaeal proteomes we found that, against the general trend for charged residues, the frequency of aspartic acid residues decreases strongly as natural growth temperature increases. In search of the explanation, we hypothesized that the reason for such unusual correlation is the deleterious consequences of spontaneous chemical transformations of aspartate at high temperatures. Our subsequent statistical analysis supported this hypothesis. First, knowing that these chemical modifications are higher in the unfolded polypeptide chains, we observed the most pronounced decrease of Asp frequency with temperature within intrinsically disordered regions. Second, it is known that the reaction rate of the chemical transformations of Asp is highest with the smallest downstream residue Gly and noticeably reduced for bulky residues. In agreement with this, the frequency of Gly residues downstream of Asp decreases with optimal growth temperature, while the frequency of bulky residues significantly increases. Thus, our analysis suggests that organisms have likely adapted to high temperatures by minimizing the harmful consequences of spontaneous chemical transformations of Asp residues.

## INTRODUCTION

Temperature is one of the most important factors that influenced the emergence of life. It is thus interesting from an evolutionary standpoint to understand the constraints that high temperature imposes on life. Furthermore, hundreds of millions of years of evolution have provided the most exhaustive of all ‘lab’ experiments to inform about how traits evolve and diversify. The reading of the traces of these experiments, in the DNA and protein sequences of extant life forms, is very powerful with the big data now available. On the genomic level, several studies explored the relationship between an optimal grow temperature (OGT) of organisms and nucleotide content of their genomes. As a result, a high G+C-content of the genomes of thermophiles (Kreil and Ouzounis, 2001) and their preference for transcription of purine-rich RNAs (Lao and Forsdyke, 2000) have been described. At the same time, most of the authors agree that thermal adaptation of proteins is especially manifest at the level of amino acid composition. Previous analyses suggested that thermophilic prokaryotes have a higher proportion of charged and bulky hydrophobic residues than mesophilic prokaryotes (Haney *et al*., 1999; Cambillau and Claverie, 2000; Kumar, Tsai and Nussinov, 2000; Perl *et al*., 2000; Fukuchi and Nishikawa, 2001; Chakravarty and Varadarajan, 2002; Zeldovich, Berezovsky and Shakhnovich, 2007). The scientific community deduced from this observation that thermophile proteins need more ionic bonds and a more densely packed hydrophobic core to stabilize their 3D structures at high temperatures (Fukuchi and Nishikawa, 2001). Our motivation for a new analysis was to confirm, refute or find new correlations between thermophilic and mesophilic proteomes by using the fast growing volume of available data.

## RESULTS AND DISCUSSION

### Frequency of aspartic acid residues decreases strongly in bacterial and archaeal proteomes as optimal growth temperature increases

To gain a handle on the big sequence data we selected prokaryotic proteomes from the BioProject database (https://www.ncbi.nlm.nih.gov/bioproject). We extracted proteomes of prokaryotic species with known optimal growth temperatures, avoiding whenever possible species with other extremophile traits (halophilia, acidophilia). To link the species data from BioProject with their proteomes from the UniProt database, we used the Taxid identifier, common to these two databases. We selected only good quality proteomes certified as “referenced proteome” by UniProt. In cases where an excess number of prokaryotic species had the same optimal growth temperature, we randomly discarded species to keep a uniform repartition of the number of organisms by temperature. For the random selection, we shuffled the candidates using a Fisher-Yates shuffle algorithm implementation from the Ruby Standard Library. This resulted in a set of 175 prokaryotes (153 bacteria and 22 archaea), growing from 16°C to 103°C. This dataset forms a resource where evenness of spread is balanced by a maximum number of proteomes (Suppl Fig. S1).

Firstly, we used our modern dataset to test the twenty-year-old proposed trends for four classes of amino acid: charged, bulky hydrophobic, hydrophilic and small amphiphilic residues, which beautifully confirms the past suggestion at high detail and probability (Fig. 1A). Indeed, the frequency of bulky hydrophobic amino acid residues (Ile, Val, Leu, Met, Phe, Tyr, Trp) in prokaryotic proteomes significantly increases with growth temperature (Fig. 1A, Spearman’s ρ = 0.57). We observed the same trend for charged residues (Lys, Arg, Glu, Asp) (Fig. 1A, ρ = 0.61). As expected, the frequency of the small amphiphilic residues, Ala, Ser and Thr, decreases with growth temperature (Fig. 1A, ρ = -0.58). These residues are frequently found in the hydrophobic cores of protein structures. Their decrease can thus be explained by the previously suggested scenario (Haney *et al*., 1999), in which bulky residues substitute for small ones and increase the packing density of protein structures in thermophiles. Other residues that decrease in frequency with growth temperature are the highly hydrophilic amino acids His, Gln, and Asn (Fig. 1A, ρ = -0.46). They are frequently located on the surface of protein structures and can be substituted with charged residues in thermophiles. The remaining Pro, Gly and Cys amino acids have very special properties and cannot be unconditionally assigned as either hydrophobic, hydrophilic or charged, amino acid residues. Their frequencies fluctuate only slightly with growth temperature. The slopes for all the individual amino acids are given in Suppl Fig. S2. We noticed that most residues conformed with their classes with the most noticeable exception being among the charged residues. Of these all of the positive correlation came from lysine and glutamate with negligible contribution from arginine and, surprisingly, an extreme antagonism from aspartate (Fig. 1B, ρ = -0.62). Revisiting previously published data based on smaller proteome datasets (Suhre and Claverie, 2003; Zeldovich, Berezovsky and Shakhnovich, 2007) we found that negative correlations of Asp with OGT were already present but has not been given due attention.

**Figure 1.**
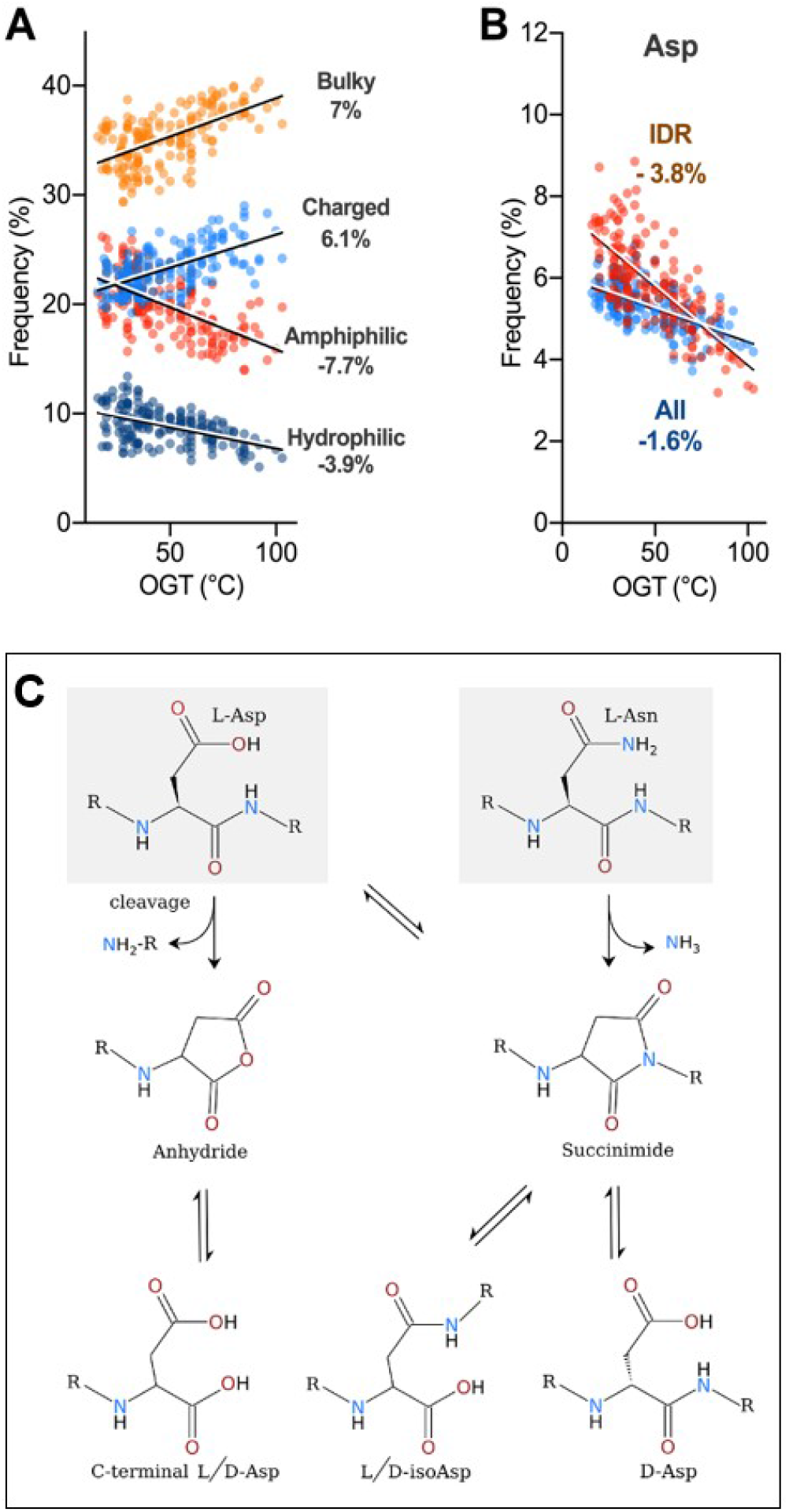
Distribution of amino acid frequencies against optimal growth temperature (OGT) in prokaryotic proteomes. **(A)** Four amino acid classes were considered: bulky hydrophobic residues (Ile, Val, Leu, Met, Phe, Tyr, Trp, orange dots), charged residues (Arg, Lys, Glu and Asp, light blue dots), small amphiphilic residues (Ala, Ser, Thr, red dots) and hydrophilic residues (His, Gln, Asn, dark blue dots). Each dot represents one proteome. **(B)** Distribution of aspartate frequency against OGT. Note aspartate bucks the trend of its charged residue group (A). Blue dots (All) represent complete proteomes and red ones (IDR) correspond to only intrinsically disordered regions of proteomes. We detected intrinsically disordered regions by using IUPred prediction (parameter ‘long disorder’) (Dosztányi et al., 2005) supplemented by a filter selecting regions of more than 16 residues with less than 25% of hydrophobic residues or with more than 20% proline. All these regions were merged and only ones of 20 residues or more were kept. Data was fitted by linear regression. Slope “a” in y=ax+b is expressed as a percent of amino acid per °C, x being expressed as °C and b as percent of amino acid. F-tests for slopes being either different or non-zero all gave p-values ≤ 10^−4^. Data analysis was done using Prism v7.0. (C) Pathways for frequently occurring spontaneous chemical reactions, which are auto-catalyzed by Asp and Asn residues. L-Asn can deamidate, form a succinimide intermediate, and then racemize (D-Asp), isomerize (iso-Asp) or form L-Asp. L-Asp can racemize (D-Asp), isomerize (iso-Asp) or auto-catalyze irreversible cleavage of the polypeptide chain via an anhydride intermediate.

### Aspartate-phobia as a reaction to harmful chemical transformations

We were intrigued to find out why Asp bucks the general trend of the charged residues, so we examined possible reasons. The observed “aspartate-phobia” of the thermophilic proteins cannot be explained by the previously described beneficial stabilization of their 3D structures by additional ionic bonds (Fukuchi and Nishikawa, 2001), because both Glu and Asp can form equally well ionic bonds with positively charged residues (Dehouck, Folch and Rooman, 2008). The other known correlations, such as a high G+C-content of the genomes of thermophiles (Kreil and Ouzounis, 2001) or their preference for transcription of purine-rich RNAs (Lao and Forsdyke, 2000) cannot explain the paucity of Asp at higher temperatures either, as aspartate codons (GAU, GAC) are not particularly G+C or purine rich.

In search of the reason, we drew our attention to the temperature-dependent chemical transformations of aspartate residue. Residues of aspartate, and the stereo-chemically similar asparagine, are both unusually prone to spontaneous chemical modifications (Collins, Waite and Van Duin, 1999). Both Asp and Asn can form a succinimide intermediate and undergo racemization (D-Asp) or isomerization (iso-Asp) (Fig. 1C). In addition, polypeptide chains can be spontaneously cleaved at Asp via an anhydride intermediate under mildly acidic pH (Partridge and Davis, 1950; Inglis, 1983) (Fig. 1C).

The large activation energies, of over 20 kcal/mol, indicate that the reaction rates towards the succinimide and anhydride intermediates and subsequent chemical transformation are highly dependent on temperature (Geiger and Clarke, 1987; Fujii, Momose and Harada, 1996). For example, increasing the temperature from 37°C to 100°C multiplied the reaction rate of deamidation by 350-fold. These chemical modifications (deamidation of Asn, racemization, isomerization of Asp and, especially, the cleavage at Asp residues) seriously and irreversibly distort or block protein functions, as shown for lysozyme (Ahern and Klibanov, 1985) and ribonuclease A (Zale and Klibanov, 1986). This is especially true for the hydrolysis of the polypeptide chain at Asp residues at high temperatures and mildly acid pH (Partridge and Davis, 1950; Inglis, 1983). Additional evidence of the biological cost of these chemical reactions is the widespread presence of the repair enzyme protein-L-isoaspartate (D-aspartate) O-methyltransferase in all three super-kingdoms of life (McFadden and Clarke, 1982). Noteworthy, this enzyme can repair spontaneous racemization, and isomerization of L-Asp, but cannot restore the hydrolysis of the polypeptide chain at Asp residues.

Increased chemical transformations of Asp with temperature likely dampen the general trend of increased charged residues with temperature and can explain the observed negative correlation of Asp frequency with growth temperature (Fig. 1B). In line with this conclusion, we observed that the percentage of Asn, which undergoes similar chemical modifications (Fig. 1C), also decreases with temperature, although with a smaller pitch (Suppl Fig. S3 ρ =-0.021). The stronger negative correlation with temperature, of Asp than Asn, can be explained by the fact that, in addition to the racemization and isomerization, only Asp not Asn can auto-hydrolyze polypeptide chains at high temperatures and mildly acid pH (Partridge and Davis, 1950; Inglis, 1983). The spontaneous and irreversible cleavage of polypeptide chain at Asp residue can thus impairs protein function more severely than other chemical modifications of Asp (isomerization and racemization) and Asn (deamination, isomerization and racemization) (Fig. 1C).

If the decreased use of Asp, and to a lesser extent that of Asn, with temperature is determined by the deleterious effects of chemical transformations, our bioinformatics analysis should also agree with two additional experimental observations. First, Geiger and Clarke, 1987 showed that spontaneous chemical modifications at Asp and Asn are higher in unfolded state of the polypeptide chains. We indeed found that the decrease of Asp and Asn frequencies with temperature is more pronounced within unfolded (intrinsically disordered, IDR) regions than in whole proteomes (Fig. 1A, 1B and Fig. S3). Secondly, Tyler-Cross and Schirch, 1991 showed that the nature of the downstream residue should modulate formation of the succinimide intermediate from Asp and Asn and of the anhydride intermediate from Asp. The reaction rate of succinimide formation is highest with the smallest downstream residue Gly and is reduced by 30 to 70-fold for bulky residues (Geiger and Clarke, 1987; Tyler-Cross and Schirch, 1991). In line with this, we found that the frequency of Gly residues downstream of both Asp and Asn residues decreases with optimal growth temperature, while the frequency of bulky residues significantly increases (Fig. 2).

**Figure 2.**
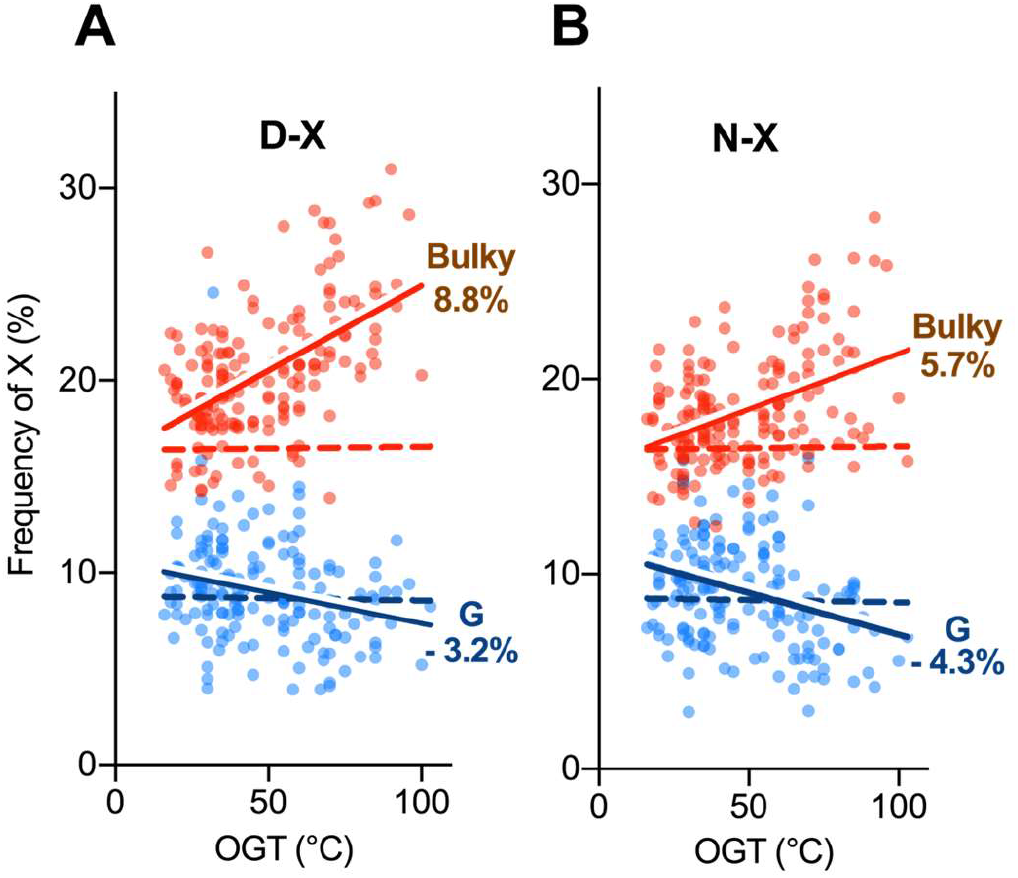
Distribution of amino acid frequencies downstream of Asp or Asn in intrinsically disordered regions of proteomes against optimal growth temperature. (panels A and B, correspondingly). X represents either bulky amino acids (Ile, Val, Leu, Met, Phe, Tyr, Trp, red dots) or glycine (Gly, blue dots). Each dot corresponds to an organism. Data were fitted by linear regression. Slopes are indicated by %. P-values of F-tests for slopes being different from the null hypothesis (dash lines) are ≤ 10^−3^ and ≤10^−4^ for G and bulky residues, respectively.

Asp and Glu are identical apart from a single extra CH_2_ extension in Glu. As thermophiles use more Glu and less Asp, the simplest adaptation to high temperatures may have been to replace Asp with Glu, which at the DNA level requires only a single transversion Y→R. Indeed, Asp to Glu is one of the most frequent replacements between mesophile and thermophile proteins (Haney *et al*., 1999).

## CONCLUSIONS

We harnessed recently increased prokaryotic proteomes to build a set of well-annotated disparate proteomes of 175 prokaryotes with optimal growth temperatures that are evenly spread from 16°C to 103°C. This allowed us demonstrate that the frequency of Asp residues is inversely correlated with optimal growth temperature. This correlation is especially surprising as it goes against the known tendency of thermophile proteins to have more ionic bonds to stabilize the 3D protein structures at high temperatures. The aspartate-phobia of the thermophilic proteins agrees with the fact that the cleavage of the polypeptide chain at Asp is the most frequent chemical reaction and has the most severe consequences for protein structure and function. The weaker negative correlation of Asn frequency with growth temperature is consistent with the lower rate of the Asn-catalyzed cleavage of proteins in comparison with the hydrolysis at Asp (Capasso *et al*., 1996, Catak *et al*., 2008). The pathway for the cleavage at Asn is indirect and requires formation of a succinimide intermediate, transformation to Asp residue and, finally, the Asp-catalyzed cleavage (Fig. 1C). The deamidation of Asn, and isomerization and racemization of either Asn or Asp, similarly to point mutations also can impair protein functions, although less efficiently than the cleavage. In agreement with chemical data that bulky residues downstream of Asx residues reduce chemical transformations, (Tyler-Cross and Schirch, 1991), the proportion of these residues downstream of Asp and Asn residues in prokaryotic protein sequences does indeed increase with the optimal growth temperature. Taken together, our observations support the notion that one of the adaptive mechanisms that thermophiles use to cope with high temperatures is to minimize the harmful consequences of spontaneous chemical transformations of Asp and Asn residues, especially, Asp-catalyzed cleavage of proteins.

Beyond this, our results can be used to build a robust model to study the environment of past lifeforms based on their inferred proteome sequences (Gaucher *et al*., 2003). Finally, in addition to more detailed insight into the molecular mechanisms of microorganisms to adapt to higher temperatures, these results offer a rational strategy for the thermo-stabilization of enzymes by protein engineering.

## Acknowledgments

The authors thank Jeremy Leclercq for help with statistical tests and generation of figures, Dr. Jean-Francois Hernandez for advice and critical reading of the manuscript and Andy Meyer for help with English. This work was supported in part by a grant to E.V. from the Ministère de l’Ėducation Nationale de la Recherche et de Technologie (MENRT).

## Author contributions

EV and AVK conceived and designed the study. EV and PF performed the analysis. AVK supervised the project. EV, PF and AVK wrote the paper.

## Data Availability

Supplementary data are available online. All other data available on request.

## References

Ahern, T. and Klibanov, A. (1985) ‘The mechanisms of irreversible enzyme inactivation at 100C’, Science, 228(4705), pp. 1280–1284. doi:10.1126/science.4001942.

Cambillau, C. and Claverie, J. M. (2000) ‘Structural and genomic correlates of hyperthermostability’, Journal of Biological Chemistry, 275(42), pp. 32383–32386. doi: 10.1074/jbc.C000497200.

Capasso, S. et al. (1996) ‘Kinetics and mechanism of the cleavage of the peptide bond next to asparagine’, Peptides, 17(6), pp. 1075–1077. doi:10.1016/0196-9781(96)00153-2.

Catak, S. et al. (2008) ‘Computational Study on Nonenzymatic Peptide Bond Cleavage at Asparagine and Aspartic Acid’, The Journal of Physical Chemistry A, 112(37), pp. 8752–8761. doi:10.1021/jp8015497.

Chakravarty, S. and Varadarajan, R. (2002) ‘Elucidation of factors responsible for enhanced thermal stability of proteins: a structural genomics based study.’, Biochemistry, 41(25), pp. 8152–61. doi:10.1021/bi025523t.

Collins, M. J., Waite, E. R. and Van Duin, A. C. T. (1999) ‘Predicting protein decomposition: The case of aspartic-acid racemization kinetics’, Philosophical Transactions of the Royal Society B: Biological Sciences, 354(1379), pp. 51–64. doi:10.1098/rstb.1999.0359.

Dehouck, Y., Folch, B. and Rooman, M. (2008) ‘Revisiting the correlation between proteins’ thermoresistance and organisms’ thermophilicity’, Protein Engineering, Design and Selection, 21(4), pp. 275–278. doi:10.1093/protein/gzn001.

Dosztányi, Z. et al. (2005) ‘The pairwise energy content estimated from amino acid composition discriminates between folded and intrinsically unstructured proteins.’, Journal of molecular biology, 347(4), pp. 827–39. doi:10.1016/j.jmb.2005.01.071.

Fujii, N., Momose, Y. and Harada, K. (1996) ‘Kinetic study of racemization of aspartyl residues in model peptides of alpha A-crystallin.’, International journal of peptide and protein research, 48(2), pp. 118–22. doi:10.1111/j.1399-3011.1996.tb00821.x.

Fukuchi, S. and Nishikawa, K. (2001) ‘Protein surface amino acid compositions distinctively differ between thermophilic and mesophilic bacteria’, Journal of Molecular Biology, 309(4), pp. 835–843. doi:10.1006/jmbi.2001.4718.

Gaucher, E. A. et al. (2003) ‘Inferring the palaeoenvironment of ancient bacteria on the basis of resurrected proteins’, Nature, 425(6955), pp. 285–288. doi:10.1038/nature01977.

Geiger, T. and Clarke, S. (1987) ‘Deamidation, isomerization, and racemization at asparaginyl and aspartyl residues in peptides. Succinimide-linked reactions that contribute to protein degradation.’, The Journal of biological chemistry, 262(2), pp. 785–94. Available at: http://www.ncbi.nlm.nih.gov/pubmed/3805008.

Haney, P. J. et al. (1999) ‘Thermal adaptation analyzed by comparison of protein sequences from mesophilic and extremely thermophilic Methanococcus species’, Proceedings of the National Academy of Sciences, 96(7), pp. 3578–3583. doi:10.1073/pnas.96.7.3578.

Inglis, A. S. (1983) ‘[28] Cleavage at aspartic acid’, in, pp. 324–332. doi:10.1016/S0076-6879(83)91030-3.

Kreil, D. P. and Ouzounis, C. A. (2001) ‘Identification of thermophilic species by the amino acid compositions deduced from their genomes.’, Nucleic acids research, 29(7), pp. 1608–15. doi:10.1093/nar/29.7.1608.

Kumar, S., Tsai, C.-J. and Nussinov, R. (2000) ‘Factors enhancing protein thermostability’, Protein Engineering, Design and Selection, 13(3), pp. 179–191. doi:10.1093/protein/13.3.179.

Lao, P. J. and Forsdyke, D. R. (2000) ‘Thermophilic bacteria strictly obey Szybalski’s transcription direction rule and politely purine-load RNAs with both adenine and guanine.’, Genome research, 10(2), pp. 228–36. doi:10.1101/gr.10.2.228.

McFadden, P. N. and Clarke, S. (1982) ‘Methylation at D-aspartyl residues in erythrocytes: possible step in the repair of aged membrane proteins.’, Proceedings of the National Academy of Sciences of the United States of America, 79(8), pp. 2460–4. doi:10.1073/pnas.79.8.2460.

Partridge, S. M. and Davis, H. F. (1950) ‘Preferential Release of Aspartic Acid During the Hydrolysis of Proteins’, Nature, 165(4185), pp. 62–63. doi:10.1038/165062a0.

Perl, D. et al. (2000) ‘Two exposed amino acid residues confer thermostability on a cold shock protein.’, Nature structural biology, 7(5), pp. 380–3. doi:10.1038/75151.

Suhre, K. and Claverie, J. M. (2003) ‘Genomic correlates of hyperthermostability, an update’, Journal of Biological Chemistry, 278(19), pp. 17198–17202. doi:10.1074/jbc.M301327200.

Tyler-Cross, R. and Schirch, V. (1991) ‘Effects of amino acid sequence, buffers, and ionic strength on the rate and mechanism of deamidation of asparagine residues in small peptides’, Journal of Biological Chemistry, 266(33), pp. 22549–22556.

Zale, S. E. and Klibanov, A. M. (1986) ‘Why does ribonuclease irreversibly inactivate at high temperatures?’, Biochemistry, 25(19), pp. 5432–5444. doi:10.1021/bi00367a014.

Zeldovich, K. B., Berezovsky, I. N. and Shakhnovich, E. I. (2007) ‘Protein and DNA Sequence Determinants of Thermophilic Adaptation’, PLoS Computational Biology. Edited by P. E. Bourne, 3(1), p. e5. doi:10.1371/journal.pcbi.0030005.

