## Supplemental Figures for "Aspartate-phobia of thermophiles as a reaction to deleterious chemical transformations"

### Supplementary data (Villain et al):

**Figure S1**.

Distribution of the proteome datasets by OGT. Zeldovich et al. (2007) dataset consists of 83 real proteomes, and three averaged proteomes at 26 °C, 30 °C, and 37 °C. Smoothing of curves was done by Prism using a window of 6 neighbouring temperatures on each side and 4th order polynomes.

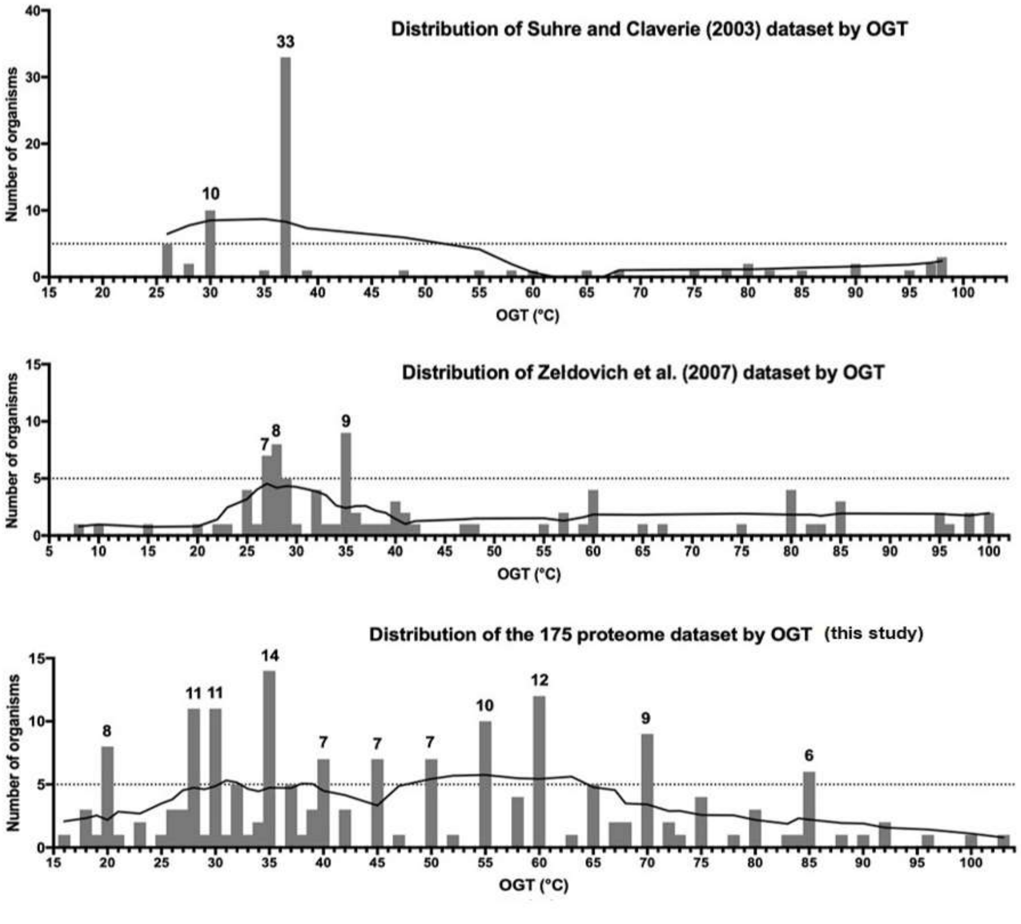

**Figure S2.**
**Distribution of the frequencies of amino acid residues in the prokaryotic proteomes depending on OGT.**

Fitting of data was obtained by linear regression. Equations are shown on top of each panel. p-values were calculated by using F-tests for slopes being non-zero. *, **, ***, **** correspond to p-values ≤ $5x10^{-2}$, ≤ $10^{-2}$, ≤ $10^{-3}$ and ≤ $10^{-4}$, respectively. Data analysis were made using Prism v7.0.

.

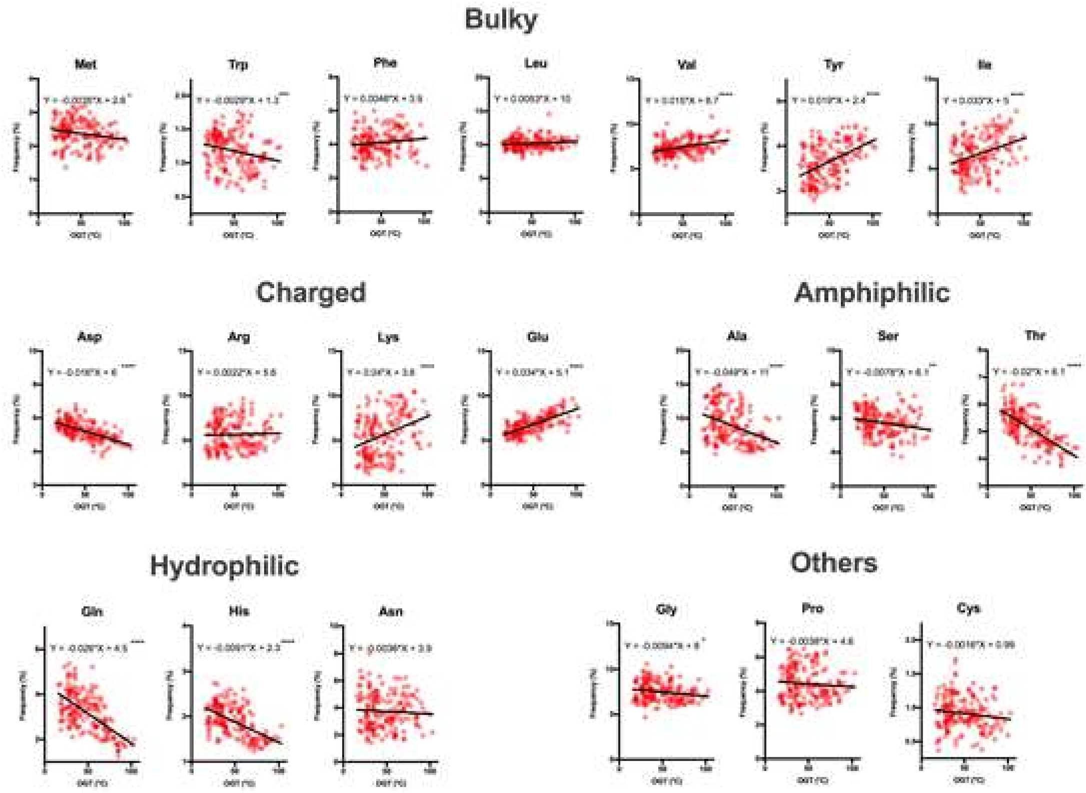

**Figure S3**. Distribution of Asn amino acid frequency against OGT. Each dot corresponds to one proteome. Blue dots (All) represent a complete proteomes and red ones (IDR) correspond to only intrinsically disordered regions of one proteomes. Fitting of data was obtained by linear regression. Slopes are indicated by %. p-values, calculated by using F-tests for slopes being either different or non-zero, all are above 0.05. Data analysis were made using Prism v7.0 .

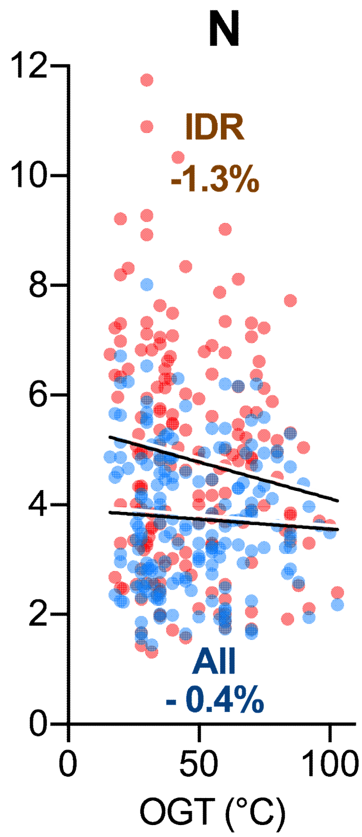
